# Pipeline to explore information on genome editing using large language models and genome editing meta-database

**DOI:** 10.1101/2024.10.16.617154

**Authors:** Takayuki Suzuki, Hidemasa Bono

## Abstract

Genome editing (GE) is widely recognized as an effective and valuable technology in life sciences research. However, certain genes are difficult to edit depending on some factors such as the type of species, sequences, and GE tools. Therefore, confirming the presence or absence of GE practices in previous publications is crucial for the effectively design and establishment of research using GE. Although the Genome Editing Meta-database (GEM: https://bonohu.hiroshima-u.ac.jp/gem/) aims to provide as comprehensive GE information as possible, it does not indicate how each registered gene is involved in GE. In this study, we developed a systematic method for extracting essential GE information using large language models from the information based on GEM and GE-related articles. This approach allows for a systematic and efficient investigation of GE information that cannot be achieved using the current GEM alone. In addition, by converting the extracted GE information into metrics, we propose a potential application of this method to prioritize genes for future research. The extracted GE information and novel GE-related scores are expected to facilitate the efficient selection of target genes for GE and support the design of research using GE.

Database Tool URLs: https://github.com/szktkyk/extract_geinfo, https://github.com/szktkyk/visualize_geinfo

## INTRODUCTION

Genome editing (GE) enables the induction of specific modifications at targeted locations in the genome^1^, promoting the advancements in genomic science. The development of Zinc-finger nuclease^2^, followed by numerous GE technologies including transcription activator-like effector nuclease (TALEN)^3^, CRISPR-Cas9^4^, base editor^5^, and prime editor^6^, has allowed for the accurate and efficient functional analysis of specific regions in genomes. GE is also applied for the transient modification of gene expression using technologies such as CRISPR activation and CRISPR inhibition^7^. Additionally, the high-throughput GE method, CRISPR screen^8^, has advanced the comprehensive investigations of genes related to specific traits using a forward genetics approach. An increase in the number of publications in which researchers use GE in their experiments^9,10^ indicates the impact of GE in life science research and how well GE technologies are accessible for application in research. Efficient and appropriate use of GE has the potential to accelerate life science research.

However, it has been reported that GE does not always work effectively at any genomics location due to various factors such as chromatin accessibility^11,12^, DNA repair mechanisms depending on the species and cell-types^13,14,15^, and target sequences^11,16,17^. These challenges arise from the fundamental mechanisms of GEs^16^. Such mechanisms include the requirement for GE tools to bind to the target sequence, the reliance on endogenous genome repair mechanisms to achieve the desired editing outcome, and the potential toxicity of the GE tools themselves. For example, TALEN has been reported to be more effective than CRISPR-Cas9 in heterochromatin regions owing to the target specificity of TALE protein’s character. In another example, a previous study revealed that 41% of GEs with CRISPR-Cas9 did not show cleavage of the targeted genome locus when they attempted to induce GE in vitro for 218 times for 153 distinct genomic loci across 18 chromosomes in the mouse genome^11^. As implied by previous studies, selecting appropriate GE tools and protocols for specific target regions is crucial to avoid repeated failures in GE experiments, especially when targeting difficult genes. In this context, access to comprehensive prior information regarding GE experiments conducted on specific genes and species serves as a valuable reference.

Several public databases provide GE-related information, including the Human Genome Editing Registry, Plant Genome Editing Database^18^, and European Sustainable Agriculture Through Genome Editing (eu-sage: https://www.eu-sage.eu/genome-search). The genome editing meta-database (GEM) is one such database, characterized by a large number of GE metadata entries from an integration of various public databases, and its focus on PubMed literature-based data collection^10^. The GEM systematically compiles and publishes metadata related to GE studies, drawing from PubMed, PubMed Central, NCBI gene, PubTator^19^, MeSH, NCBI taxonomy, and EXTRACT2.0^20,21^, integrating these public databases and services to provide as comprehensive data as possible. The GEM allows users to search for 46,039 GE-related articles as of September 18, 2024, linked with associated GE tools, genes, species, cell types, mutation types, diseases, BioProject IDs, and Gene Expression Omnibus (GEO) IDs. As each article may be linked to a few genes, the current GEM as of September 18, 2024, contained 92,182 entries in total, with the data visualized according to GE tools, species categories, and years. Such a comprehensive database is useful for exploring various research scenarios, including identifying genes that are accessible or efficiently targeted by GE, genes that remain understudied in the GE research field (i.e., genes that have not been studied or paid attention to), and GE cases involving orthologs of the gene of interest. However, the GEM is currently in the prototype stage, and several issues limit its use for these purposes. One issue is that all collected genes linked to GE articles in GEM are displayed without indicating their specific role because of the data collection methodology used for GEM. As a result, it is unclear from the current GEM whether a gene was the target of GE, if it was a gene that reported as altered expression due to GE of other genes, or if the entry was a result of a data collection error.

In this study, we identified one of the metadata issues in the GEM and proposed a novel pipeline to improve metadata by addressing this issue. First, we analyzed a subset of the GEM dataset to clarify this issue. By analyzing 266 annotation pairs of gene ID and PubMed ID as a case study, we identified four different types of genes in the GEM. This makes it difficult for users to search for specific GE-related genes. To address this issue, in which the current GEM does not specify the role of each gene in GE, we developed a method that utilizes large language models (LLMs). LLMs, such as OpenAI’s ChatGPT and Meta’s Llama, have been reported to exhibit strong capacities for a wide range of complex tasks beyond text generation^22^, including the extraction of metadata from research literature^23–25^. For example, LLMs fine-tuned on a relatively small dataset of 400-650 annotated text-extraction pairs can achieve F1 scores above 0.8 in named entity recognition and relation extraction (NERRE) tasks from scientific texts^23^. Previous studies have demonstrated the effectiveness of LLMs in both entity recognition and relation extraction^23,24^. Furthermore, even without fine-tuning, prompt engineering and conversational follow-up processing have been shown to enable highly accurate metadata extraction^25^. Therefore, we selected LLMs to address the challenges posed in this study. The pipeline developed in this study (Figure 1) allows users to access genes targeted by specific species and GE tools more accurately than the current GEM dataset. Additionally, it enables users to roughly estimate the number of cases “a gene was targeted by GE (GE_target_gene)” and the number of articles “a gene reported as altered expression due to GE of other genes” (GE_deg)”. This study proposes a novel methodology to explore GE information through a systematic and comprehensive PubMed literature-based analysis facilitated by a combination of GEM data and LLM.

**Figure 1.**
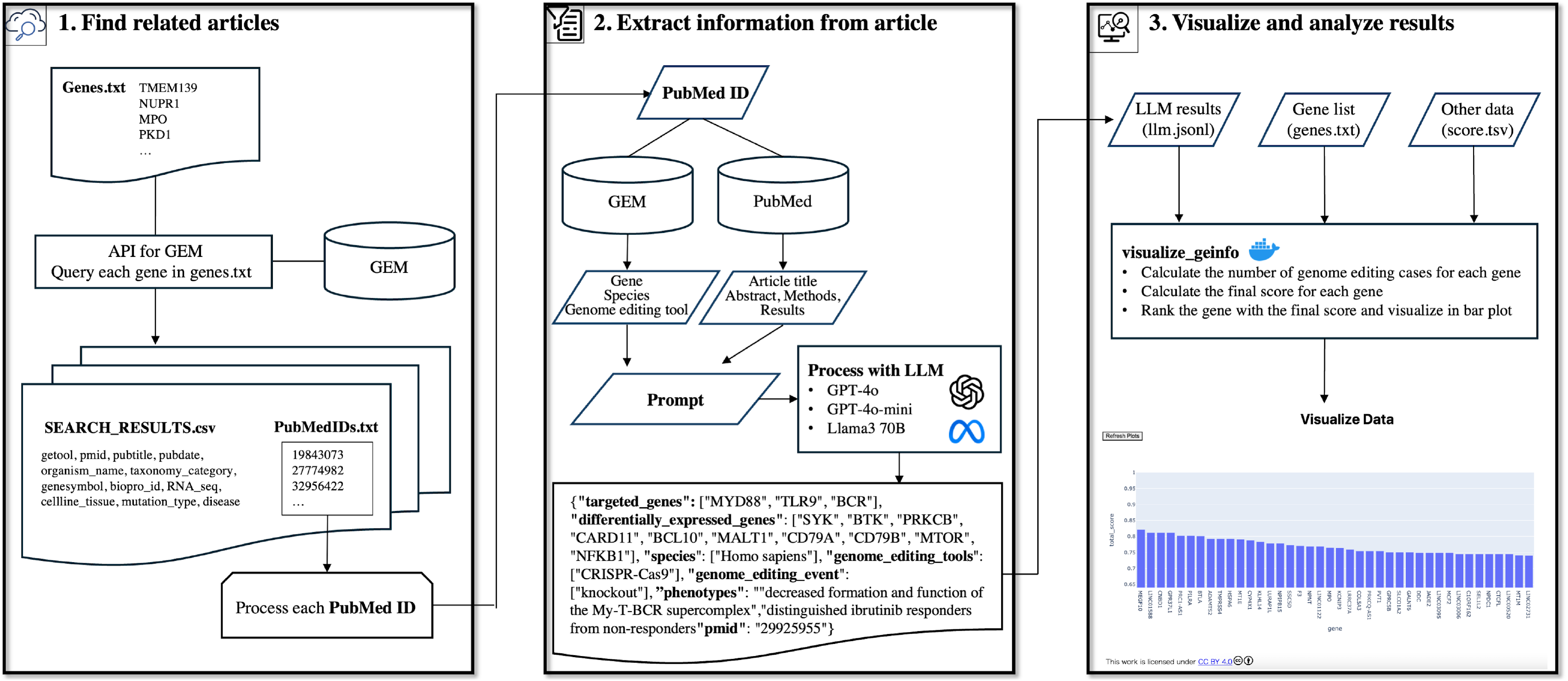
Overview of the pipeline for exploring information on genome editing using the genome editing meta-database and a large language model. The pipeline comprised three major steps: Step 1 retrieves genome editing-related publications for LLM processing. The system queries GEM to obtain genome editing-related publications associated with genes of interest. Step 2 implements LLM-based information extraction, utilizing multiple inputs including GEM’s gene annotations, GEM’s species information, GEM’s genome editing tool information, literature text, and prompts to extract detailed genome editing information. Step 3 facilitates the visualization of extracted data and implements scoring metrics.

## MATERIALS AND METHODS

### Overview of the pipeline

The pipeline developed in this study systematically collects novel metadata related to GE by utilizing the GEM and literature as reference materials processed through LLM. Figure 1 shows an overview of the entire pipeline. In Step 1, “Find related articles” selects relevant GE articles for subsequent analysis based on a group of gene IDs or symbols of interest. In Step 2, “Extract information from article”, applies LLM-based processing to the selected articles to extract GE-related information. In Step 3, “Visualize and analyze results,” visualizes and analyzes the extracted information (Figure 2). A detailed explanation of each step is provided in the subsequent sections.

**Figure 2.**
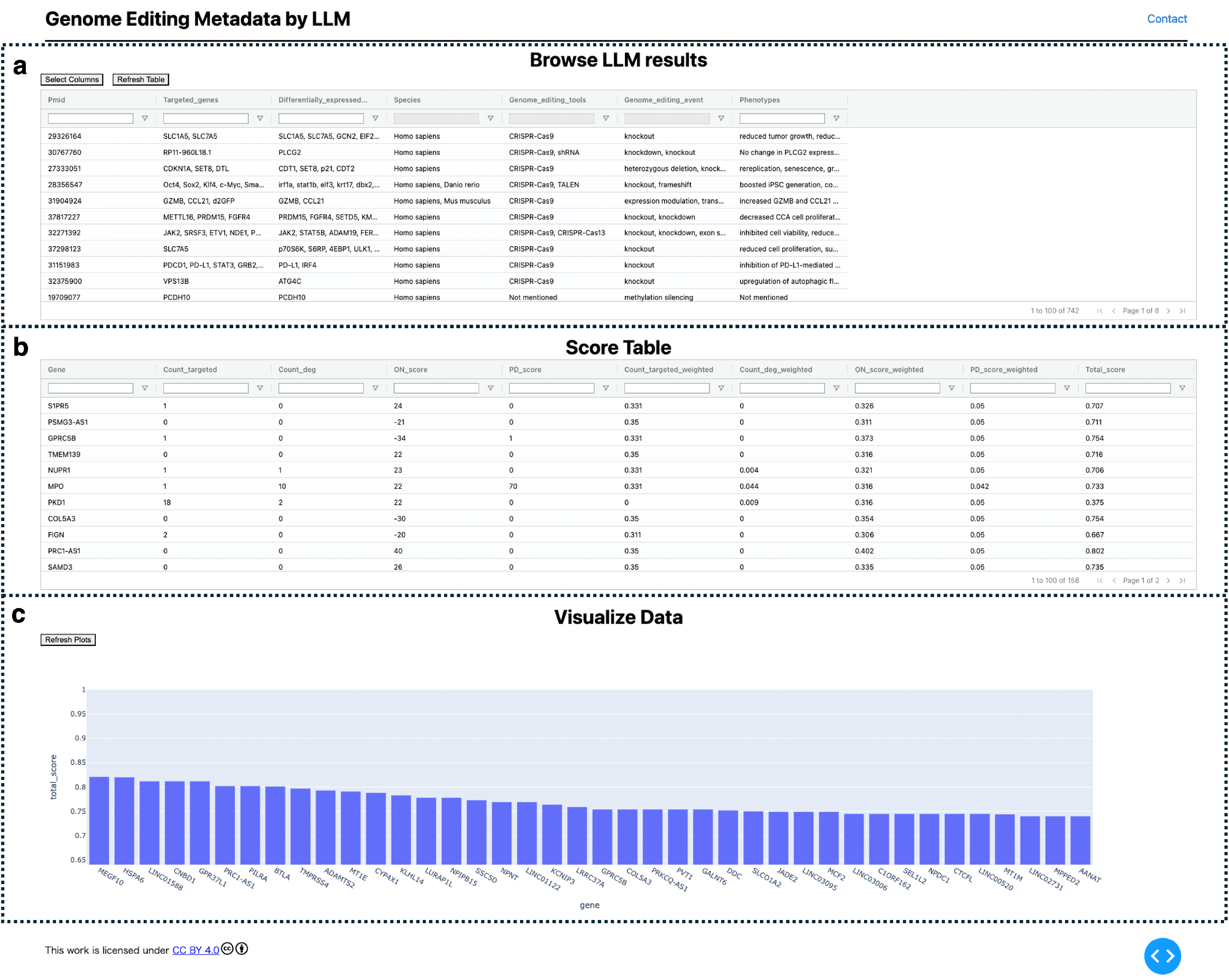
Visualization of extracted genome editing information. (a) Extracted information provided in a searchable table. (b) genome editing related metrics, and additional custom scores are provided as a table. In addition, the normalized scores were calculated and are listed in table. (c) The top 40 ranked genes are visualized using a bar plot with their scores.

### Step 1: Find related articles

We conducted a case study focusing on the GEM entries related to the 168 genes (See Table S1 in the Supplementary File). These 168 genes represented by Ensembl IDs were identified in a previous meta-analysis conducted by our group as being associated with oxidative stress in Parkinson’s disease^26^. Each GEM entry comprises a GE-related article linked to specific genes and other associated metadata. To retrieve GEM entries for the 168 genes, we used the “gem_api” (https://github.com/szktkyk/gem_api) to query the GEM dataset. A text file containing the NCBI Gene IDs or gene symbols, separated by line breaks, was prepared, and the script “01_search_gem.py” from the “extract_geinfo” repository (https://github.com/szktkyk/extract_geinfo) was executed to retrieve relevant GEM entries. As GEM only accepts NCBI Gene ID (to search for a species-specific GEM entries) or gene symbol (to search for orthologs across species), we converted the 168 Ensembl IDs to NCBI Gene IDs and gene symbols. Out of the 168 genes, 146 had an associated NCBI Gene ID, and 158 had a gene symbol as a result of gene ID conversion using NCBI command-line tools (https://www.ncbi.nlm.nih.gov/datasets/docs/v2/download-and-install/) (See Table S1 in the Supplementary File).

### Step 2: Extract information from article

Using the custom-developed scripts “extract_geinfo” (https://github.com/szktkyk/extract_geinfo), we performed information extraction using LLMs. We have conducted the experiment of Step 2 twice for the following reasons: 1) To explore and optimize our choice of LLMs, 2) To explore and optimize the prompt, 3) To explore the genome editing information we should extract, and 4) To enable two separate rounds of performance evaluation. For the first round of experiments in May 2024, we have processed 259 GE-related articles retrieved in Step 1 (human-specific GEM entries retrieved by querying with 146 NCBI Gene IDs). For the second round of experiments in August 2024, we have processed 742 GE-related articles retrieved in Step 1 (GEM entries across species retrieved by querying 158 gene symbols). For each article, a specific prompt was created incorporating reference information, including metadata from the GEM entries and textual data from the article. This process was repeated 259 or 742 times, corresponding to the number of articles, to enable comprehensive information extraction from the GE-related articles associated with the queried genes archived in the GEM. Specifically, we prepared a list of PubMed Central IDs (PMCIDs) with text file. By running the “02_get_pubdetails.py” script from the extract_geinfo repository, we collected detailed information such as the textual content of each article. Subsequently, executing “03_run_gpt.py” facilitated information extraction from the textual data of PubMed Central articles by referencing the metadata from GEM entries.

Two different prompts were tested for Step 2. In the first round of experiments, the following prompt was used to extract eight types of metadata: “targeted genes of GE,” “genes reported as altered expression due to GE of other target genes,” “Species studied using GE,” “GE tools used,” “GE events induced,” “Study context,” “Key findings,” and “Implications suggested.”

In the second round of experiments, we refined the prompt as follows and extracted six types of metadata: “targeted genes of GE,” “genes reported as altered expression due to GE of other target genes,” “Species studied using GE,” “GE tools used,” “GE events induced” and “Phenotypes resulting from GE.”

In the first round of experiments, we utilized the following information: the genes and species linked to the GEM entries as well as the titles and abstracts of the articles. In the second round of experiments, we incorporated additional information, including the GE tools registered in the GEM entries and the textual content from the methods and results sections of the article.

The outputs from the LLM were standardized into a unified JSON format using the “output_parser” function from the LangChain library. Four LLMs were tested: GPT-4, GPT-4o, GPT-4o-mini, and Llama3-70b. In the first round of experiments, each information extraction task was performed once using GPT-4, GPT-4o, and Llama3-70b models. In the second round, each task was performed once using GPT-4o and GPT-4o-mini models. Based on its accuracy, GPT-4o was selected for further use. The primary libraries and tools used to query the LLM API were LangChain (https://www.langchain.com/) and Groq (https://pypi.org/project/groq/).

### Step 3: Visualize and analyze results

The information extraction results using LLM and GEM were visualized locally as a table_vis1 (Figure 2a) using the custom-developed tool “visualize_geinfo” (https://github.com/szktkyk/visualize_geinfo), which allows for automated calculation of GE related metrics and facilitates easy searching of the results. During visualization, the number of GE targeted cases (GE_target_count) and the number of articles reporting gene expression changes due to GE of other genes (GE_degs_count) for each of the queried genes (158 genes in this case) were automatically calculated and displayed as table_vis2 (Figure 2b). “visualize_geinfo” tool has built in functionality to incorporate additional custom-related metrics in table_vis2 (Figure 2b). In this study, we added two metrics: scores from a transcriptomics meta-analysis (meta-analysis score), and the number of articles reporting an association with Parkinson’s disease for each gene (PD_count)^26^ (Figure 2b). Meta-analysis score was calculated in the previous study^26^. This score represents the number of RNA-seq data pairs showing expression changes out of data pairs derived from 10 research projects, suggesting the gene’s association with oxidative stress. A higher absolute value of the score indicates more consistent differential expression across multiple studies, suggesting the gene’s association with oxidative stress. We also applied max-min normalization to scale each value between 0 and 1 (equation1), and calculated a cumulative score, which is presented in table_vis2 (Figure 2b). In total, 158 genes were ranked in descending order based on cumulative scores. For metrics in which lower values indicated stronger target candidates (i.e., GE_target_count and PD_count), the scoring system adjusted the ranking by subtracting each value from the overall maximum (equation2). The score is automatically calculated through executing the “weight_score.py” in visualize_geinfo repository. Two important settings - addition of custom-related scores to the table_vis2 and indication of specific metrics where lower values indicating stronger target candidates, can be configured by editing the “config.py” file in the repository. The top 40 ranked genes were visualized using a bar plot (Figure 2c). The visualize_geinfo tool can be executed in a containerized environment by loading a Docker file from the repository, ensuring that it can run on any computer platform.

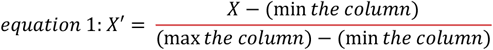

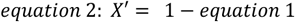

### Preparation of evaluation data

After querying 146 NCBI Gene IDs against the GEM dataset of 7 May 2024, 266 annotation pairs between the NCBI Gene ID and PubMed ID were retrieved from 259 articles. We manually curated the role of each gene from 266 annotation pairs by reviewing the contents of the annotated articles. The curation results are summarized in the Table S2 of the Supplementary File. If the gene was identified as a target of GE in the article, “1” was entered in the “curation_gene” column of the curation file. If the gene was not a target, “0” was recorded. For genes targeted in non-human species, “2” was entered, with the correct species was noted in the memo column.

If the gene was reported in the article to have altered expression due to GE of other genes, “1” was entered in the “deg” column of the curation file, and “0” if not. If a gene was already labeled as a GE target, “2” was entered in the deg column to exclude it from LLM performance evaluation. This exclusion was necessary because LLMs might infer expression changes in genome-edited genes as GE_deg, even without explicit mention in literature.

### Performance evaluation of information extraction by large language model

The evaluation of LLM-based information extraction was conducted for two types of extracted metadata: genes targeted by GE (GE_target) and genes reported as having altered expression due to the GE of other genes (GE_deg).

For GE_target, the evaluation followed these criteria: if the evaluation data (curated csv file) was labeled “1” and the LLM output included the gene, it was classified as a true positive (TP); if the gene was missing from the LLM output, it was a false negative (FN). If the evaluation data was labeled “0” and the LLM output did not include the gene, it was considered a true negative (TN), while inclusion of the gene was treated as a false positive (FP). For cases labeled “2,” it was considered a TP if the gene was present in the LLM output and the species mentioned in the LLM results matched the curated species in memo column.

For GE_deg, 163 annotation pairs were evaluated, excluding the 103 annotation pairs curated as GE_target. The same criteria used for GE_target applied: if the evaluation data was labeled “1” and the gene was included in the LLM output, it was classified a TP, while omission of the gene was an FN. For cases labeled “0,” if the gene was absent from the LLM output, it was considered a TN; otherwise, it was classified as an FP.

The accuracy, precision, recall, and F1 scores were calculated using the following formulas:

Accuracy = (TP+TN)/(TP+TN+FP+FN)

Precision = TP/(TP+FP) Recall = TP/(TP+FN)

F1 score = 2*(Precision*recall)/(Precision+Recall)

The calculation was executed by running “evaluate_targetedgenes.py” and “evaluate_deg.py” from extract_geinfo repository (https://github.com/szktkyk/extract_geinfo).

## RESULTS

### Proportion of genome editing targeted genes in Genome Editing Meta-database

As mentioned in the introduction, the Genome Editing Meta-database (GEM) dataset includes various types of genes linked to GE related literature owing to the nature of its data collection system. To investigate the types of genes linked to GE-related article, we manually curated a subset of the GEM dataset. Following Step 1 of the pipeline (Figure 1, method step1), we queried 146 NCBI Gene IDs against the GEM dataset to retrieve GE-related articles. These 146 NCBI Gene IDs were converted from 168 Ensembl Gene IDs that were previously identified through data-driven analyses as responsive to oxidative stress in Parkinson’s disease^26^. Consequently, we retrieved 266 unique pairs of annotations, where each pair contained a gene ID (NCBI gene ID) matched with its corresponding article ID (PubMed ID). As some articles were linked to multiple genes, the total number of GE-related articles retrieved was 259 (referred to as 259_GE_articles).

Manual curation of the 266_GE_annotation pairs using the method outlined in preparation of evaluation data, revealed four major categories of genes: genes targeted by GE (38.72%, 103 of 266_GE_annotation_pairs), genes reported as altered expression due to GE of other genes (24.44%, 65 of 266_GE_annotation_pairs), genes studied in the article but not related to GE (10.53%, 28 of 266_GE_annotation_pairs), and genes collected due to extraction errors (8.27%, 22 of 266_GE_annotation_pairs).

If these results reflect a similar trend across the entire GEM dataset, which contains 92,182 entries as of September 18, 2024, this suggests that only approximately 39% of the search results represent the intended entries if a user searches for GEM to investigate whether a gene has been targeted by GE. Additionally, approximately 19% of the search hits would consist of unrelated studies or extraction errors. This indicates that the GEM dataset, in its current form, may be an unreliable dataset for such purposes.

### Proportion of genome editing targeted genes after information extraction by large language model

To address the challenge of identifying the role of each gene in a GE study, we explored the application of large language models (LLMs). In Step 2 of the pipeline, we utilized information on genes, species, GE tools from the GEM, along with textual data from the articles, combined with custom prompts, as input to the LLMs to systematically extract GE metadata.

In May 2024, as part of the first round of experiments, we used genes and species data from the GEM, and the titles and abstracts of relevant articles. Several LLMs were tested to extract “targeted genes by GE” and their extraction performance, cost, and processing time were compared. As outlined in the ‘Materials and Methods: Step 2: Extract information from the article’, 259_GE_articles were processed using LLMs, and the extracted information was compared with manually curated evaluation data (See Table S2 in the Supplementary File). The results show that GPT-4 achieved 84.96% accuracy, a cost of $9.9, and a processing time of 3000 seconds; GPT-4o achieved 90.23% accuracy, a cost of $1.4, and a processing time of 960 seconds; and Llama3 70B achieved 83.83% accuracy, a cost of $1.5 (the cost of using GPT-4 with LangChain’s output_parser library), and a processing time of 4,440 seconds (Table 3). Llama3-70B was tested using the Groq’s API Demo version. Based on these results, GPT-4o, which had the highest accuracy and a relatively low cost, was selected.

**TABLE 1:**
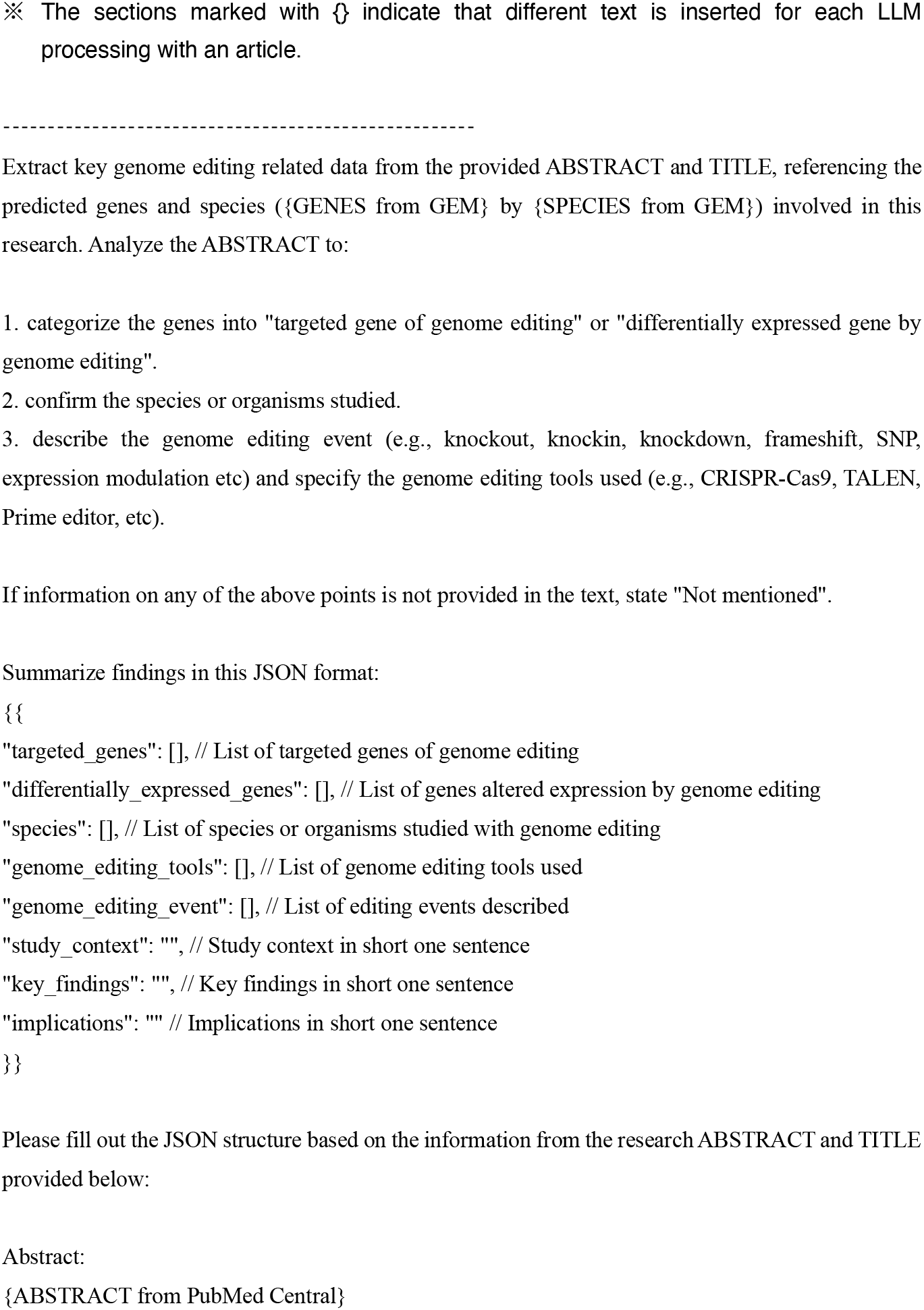
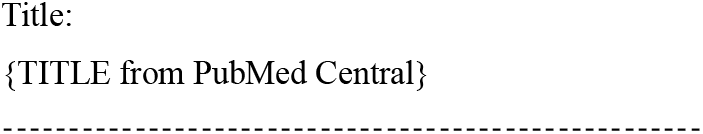
the prompt used for the first round of genome editing information extraction.

**TABLE 2:**
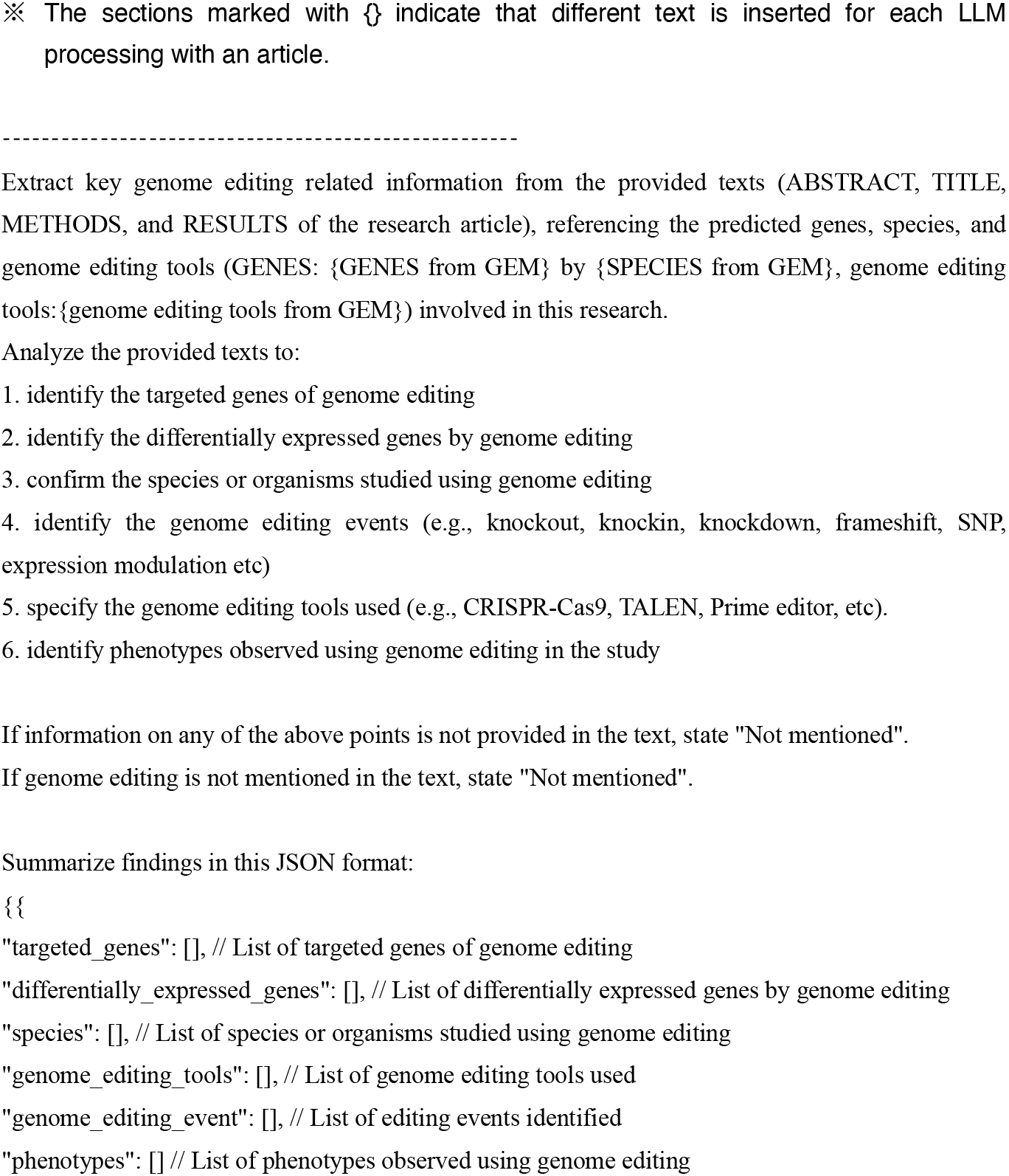
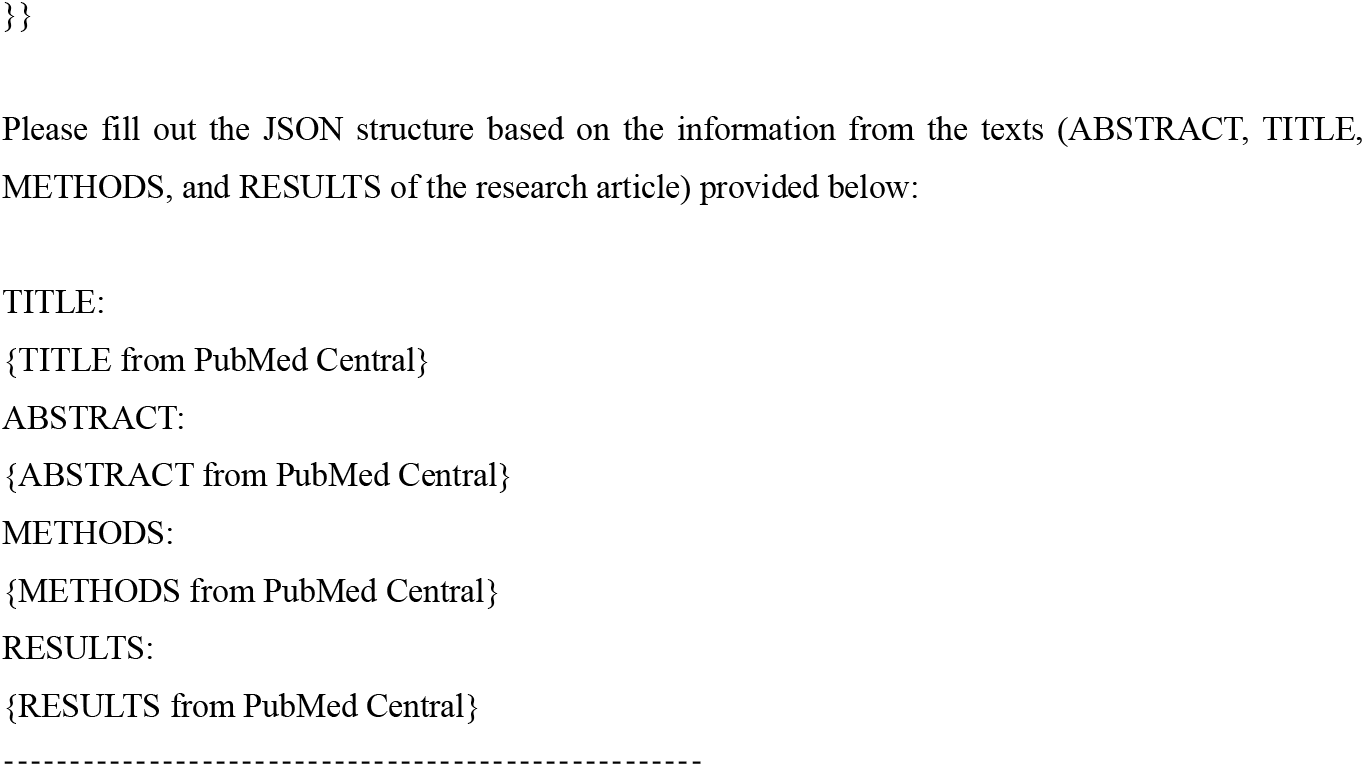
the prompt used for the second round of genome editing information extraction.

**TABLE 3:** Evaluation for extraction of GE_target (genes targeted by GE) using large language models for the first round of experiments (20240510-16)

|  | gpt-4 | gpt-4o | llama3-70b |
| --- | --- | --- | --- |
| number of articles processed | 259 | 259 | 259 |
| accuracy for 259 articles | 0.8496 | 0.9023 | 0.8383 |
| precision for 259 articles | 0.7890 | 0.9326 | 0.7679 |
| recall for 259 articles | 0.8349 | 0.8058 | 0.8350 |
| f1 for 259 articles | 0.8113 | 0.8646 | 0.8000 |
| price | \$9.90 | \$1.40 | \$1.50 |
| time | 3000 (s) | 960 (s) | 4440 (s) |

In August 2024, during the second round of experiments, the prompt was refined to improve accuracy, and with the release of GPT-4o-mini, we conducted a comparison between GPT-4o and GPT-4o-mini. In the second round of experiments, our custom prompts for LLMs included information about genes, species, and GE tools from the GEM, and textual data from the title, abstract, methods, and results sections of the article. Metadata extraction was performed using GPT-4o and GPT-4o-mini to extract six types of metadata: targeted_genes (genes targeted by GE, referred to GE_target), differentially_expressed_genes (genes reported as altered expression due to GE of other genes, referred to GE_deg), species (organisms studied using GE), genome_editing_tools (tools used for GE in the study), genome_editing_event (events induced by GE), and phenotypes (phenotypes observed as a result of GE). As listed in Table 4, GPT-4o outperformed GPT-4o-mini and the models from the first round of experiments in terms of extraction accuracy. Therefore, the results of GPT-4o on the second round were selected to use for further analysis.

**TABLE 4:** Evaluation for extraction of GE_target and GE_deg (genes reported as altered expression due to GE of other genes) using GPT-4o for the second round of experiments (20240802)

|  | gpt-4o-mini (GE_targeted) | gpt-4o (GE_targeted) | gpt-4o (GE_deg) |
| --- | --- | --- | --- |
| number of articles processed | 742 | 742 | 742 |
| accuracy for 259 articles | 0.8797 | 0.9511 | 0.8528 |
| precision for 259 articles | 0.7840 | 0.9167 | 0.7662 |
| recall for 259 articles | 0.9615 | 0.9612 | 0.9077 |
| f1 for 259 articles | 0.8596 | 0.9384 | 0.8310 |
| price | \$0.76 | \$26.55 | \$26.55 |
| time | 2050 (s) | 4339 (s) | 4339 (s) |

For GE_target, the results from the second round of experiments demonstrated that the LLM was able to interpret the context of GE-related genes described in the article, with an F1 score of 0.9384. For GE_deg, the F1 score was 0.831.

In the task of extracting GE_target from the article and GEM data using LLM, precision had the lowest value among the measured metrics. Of the 266_GE_annotation_pairs, 13 genes were incorrectly extracted, including nine false positives (FP) and four false negatives (FN) (Table 5). Six of the nine FP cases resulted from the LLM incorrectly annotating GE_deg as GE_target. Similarly, precision was the lowest metric in the GE_deg extraction task with 19 FP and six FN cases (Table 6). In the evaluation of the two types of GE-related metadata in this study, a trend was observed in which precision was lower compared to accuracy and recall (Tables 5 and 6). These results suggest that LLMs tend to over-interpret contextual information rather than overlook it, particularly when distinguishing between closely related concepts such as GE_target and GE_deg. This tendency might be attributed to the LLM’s inherent behavior of making inferences beyond explicitly stated information.

**TABLE 5:**
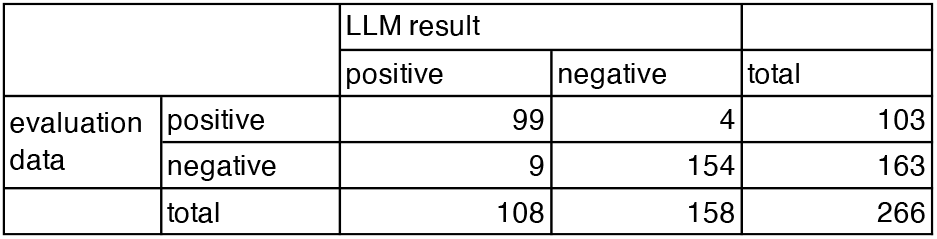
Confusion matrix for the GE_target of the second round of experiments with GPT-4o (20240802)

|  |  | LLM result |  | total |
| --- | --- | --- | --- | --- |
|  |  | positive | negative |  |
| evaluation data | positive | 99 | 4 | 103 |
|  | negative | 9 | 154 | 163 |
|  | total | 108 | 158 | 266 |

**TABLE 6:** Confusion matrix for the GE_deg results of the second round of experiments with GPT-4o (20240802)

|  |  | LLM result |  | total |
| --- | --- | --- | --- | --- |
|  |  | positive | negative |  |
| evaluation data | positive | 59 | 6 | 65 |
|  | negative | 18 | 80 | 98 |
|  | total | 77 | 86 | 163 |

The output files include the results of the GPT-4o and GPT-4o-mini for the second round of experiments (See Tables S3 and S4 in the Supplementary File). The evaluation results for GE_target and GE_deg for the second round of experiments using GPT-4o are available in the Tables S5 and S6 Supplementary File.

### Calculation of genome editing-related metrics for genes

By utilizing an LLM-based information extraction process, we were systematically able to obtain novel GE metadata that the current GEM cannot achieve. Although the evaluation was conducted on only 259_GE_articles, we expanded the dataset to include 742 articles retrieved by querying the GEM with 158 gene symbols rather than NCBI Gene IDs. The searching for gene symbols in the GEM enables the inclusion of articles linked to orthologous genes in non-human species, resulting in an increased number of hits. We processed these 742 articles following Step 2 of the pipeline to collect GE metadata. From the LLM results for these 742 articles, two GE-related metrics were calculated for each gene: (1) the number of cases in which the gene was targeted by the GE tool (GE_target_count), and (2) the number of articles reporting altered gene expression due to GE of other genes (GE_deg_count). These metrics were automatically calculated by running the custom-developed visualize_geinfo tool (https://github.com/szktkyk/visualize_geinfo), which calls search_gem_llm.py to compute how frequently each gene was identified as “GE_target_count” or “GE_deg_count” based on the LLM output.

The top 10 genes with the highest GE_target_count values among the 158 genes were: *PKD1, FOS, SLC7A5, TXNIP, STAT6, SPP1, SLC3A2, TUBB3, SLC26A4*, and *CARD11*. These genes are considered to have a higher likelihood of being well-studied for effective GE within the 158 gene set. In contract, 103 of the 158 genes had a GE_target_count of zero according to the GEM and LLM, suggesting that they have not yet been extensively studied using GE or have limited examples. These are challenging candidate genes, potentially leading to novel discoveries through GE experiments.

The top 10 genes with the highest GE_deg_count values among the 158 genes were: *FOS, SPP1, TXNIP, MPO, CDKN1C, CRYAB, CD74, STAT6, CPT1A*, and *PKD1*. These genes have been suggested to have potential responsiveness or exhibit certain phenotypes in response to GE.

### Utilization of genome editing-related metrics

As an experimental approach for utilizing the two GE related metrics, we scored and ranked the 158 genes. In ranking the 158 genes, we incorporated two custom metrics: a meta-analysis score derived from gene expression data (meta-analysis_score) and the number of studies reporting an association with Parkinson’s disease (PD_score), along with two GE-related metrics: GE_target_count and GE_deg_count. The custom-developed visualize_geinfo tool allows the addition of custom scores (meta-analysis_score and PD_score) to the default metrics (GE_target_count and GE_deg_count) in the table. All four metrics were normalized to a 0-1 scale using max-min normalization with weights assigned based on importance. These weights are configurable through the config.py file, allowing users to adjust them according to their priorities when running Step 3 of the pipeline. In this study, we assigned a weight to each metric as follows: GE_target_count:0.35, GE_deg_count:0.15, meta-analysis_score:0.45, and PD_score:0.05. we assigned the highest weight (0.45) to the meta-analysis_score as it represents results from transcriptome meta-analysis, which indicates the functional significance of genes in terms of expression changes. The second-highest weight was given to GE_target_count, as we aimed to prioritize genes that had not been previously studied using genome editing. GE_deg_count received the third-highest weight, as it potentially indicates gene functionality from an expression perspective. While PD_score, which reflects the research attention a gene has received in Parkinson’s disease studies, may be significant in certain contexts, it was given the lowest priority in this study.

After calculating the total score for each gene, the top 40 genes were ranked and visualized in a bar plot (Figure 2c). The table shown in Figure 2b displays all 158 genes sorted by score. The top 10 ranked genes were *MEGF10, HSPA6, LINC01588, CNBD1, GPR37L1, PRC1-AS1, PILRA, BTLA, TMPRSS4*, and *ADAMTS2*. The scores for the 158 genes are listed in the Table S7 of the Supplementary File. We also examined the top-ranked genes when the weights were slightly changed as follows: GE_target_count:0.25, GE_deg_count:0.10, meta-analysis_score:0.55, and PD_score:0.10. We observed that the composition of the top 10 genes remained consistent except for a swap between the first and second positions. The scores for the 158 genes with these modified weights are listed in Table S8 of the Supplementary File.

These top-ranked genes can be hypothesized as potential targets for future research, as they scored highly in the meta-analysis_score based on gene expression meta-analysis and the GE_deg_count based on literature interpretation by the LLM. In addition, they have low GE_target_count and PD_score values, indicating that they are understudied in the GE and PD research fields. This suggests that they may lead to new discoveries on in-depth investigations. However, these top-ranked genes have not yet been experimentally validated.

## DISCUSSIONS

This study identified a key issue with the existing genome editing meta-database (GEM), in which genes linked to GE-related literature in the GEM can be classified into four major categories; however, the current GEM does not allow for determining which category each gene belongs to. To address this limitation, we employed large language models (LLMs). In a case study involving 259_GE_articles associated with the test 168 genes (including 146 NCBI Gene IDs and 158 Gene Symbols), we extracted information on GE contexts, including targeted genes of GE (GE_target) and genes reported as having altered expression due to GE of other genes (GE_deg). The results showed that GE_target was identified with an F1 score of 0.9384, and GE_deg with an F1 score of 0.8310. This demonstrates that combining LLMs with existing GEM can provide novel GE metadata that could not be systematically collected previously. By leveraging this enhanced GE information, we calculated two new metrics for each gene: “number of GE targeted cases (GE_target_count)” and “number of articles reporting differential expression due to GE of other genes (GE_deg_count).” These metrics were then used in an experimental approach to score and prioritize the 158 genes.

After normalizing the scores, we ranked the 158 genes and examined the characteristics of some of the top-ranked genes. For example, *LINC01588*, ranked third, had a score of zero for GE_target_count, GE_deg_count, and PD_score. Its high ranking was driven by its relatively strong oxidative stress (OS) meta-analysis_score. A PubMed search for “LINC01588” returned five articles, one of which reported that knocking down *LINC01588* enhances oxidative stress through its interaction with HNRNPL^27^. *LINC01588* encodes for a non-coding RNA, which may explain the limited research on this gene. The second-ranked gene, *HSPA6*, encodes the molecular chaperone heat shock 70kDa Protein 6, which responds to stress. It had three cases of GE, and one study linking to PD. With the value of 2 for GE_deg_count and a meta-analysis_score of 50, *HSPA6* ranked in the second position. As one of the GE articles also explained *HSPA6* as an upregulated gene in PD^28^, *HSPA6* is an already established target gene for PD and can be considered a positive control for this ranking. *CNBD1*, ranked fourth, was downregulated in OS conditions. The protein encoded by cyclic nucleotide-binding domain-containing 1 remains relatively understudied, with limited available information. Research on *CNBD1* in PD or OS is sparse, no GE cases were found using our method, and no PubMed hits querying with “*CNBD1*” were identified, making *CNBD1* as one of the potential new research target gene for the future PD or OS study.

A high score on these metrics does not necessarily imply that the genes are functionally linked to OS or PD. However, by incorporating GE information, this method can be seen as a way to prioritize candidate genes for further experimental functional analysis, particularly focusing on genes with limited prior research, but notable gene expression profiles. There is a growing concern regarding research bias toward well-studied human genes^29,30^. In response, approaches aimed at promoting the investigation of understudied genes and fostering new discoveries, such as omics-based methods as “Unknomics”, have gained attention^31–33^. Unknomics seeks to advance research on novel genes. This scoring method, which integrates GE_target_count and PD_score in this study, can be viewed as a form of unknomics.

In the end, we outline the four major limitations of the pipeline developed in this study. First, the GEM dataset was collected based only on the literature from PubMed. As of September 18, 2024, the GEM contained 46,039 GE-related articles retrieved from PubMed using a custom search query (https://github.com/szktkyk/gem/blob/main/config.py). However, there is a possibility that irrelevant literature unrelated to GE was included, and relevant literatures may have been missed. Articles that were not indexed in PubMed were excluded from the GEM dataset. Second, the number of articles processed using LLMs was limited in this study. As a feasibility study, only 742 of the 46,039 articles in the GEM dataset were processed using the pipeline. The evaluation data were limited to 259 of the 46,039 articles, indicating that the performance of the LLM was assessed only on a subset of the data registered in the GEM. The LLM performance on a large scale remains untested. Third, our evaluation of GE metadata was limited to GE_target and GE_deg due to insufficient labeled data for other metadata types. Additionally, we have not assessed how variations in extracted metadata types between the first and second rounds might affect the extraction performance of GE_target and GE_deg. Finally, there is always a risk of error in the LLM output. Information extraction for GE_target and GE_deg using LLM achieved accuracies of 95.11 and 85.28%, respectively, indicating error rates of 4.89 and 14.72% for GE_target and GE_deg. Thus, potential errors in the LLM output cannot be entirely ruled out. Although this pipeline shows the potential for systematically collecting a large amount of GE metadata for future research, the results generated by the LLM should not be regarded as definitive.

## CONCLUSION

In this study, we explored a method to address the metadata issues in GEM using LLMs. Through this approach, we were able to systematically collect metadata that could not be obtained through the conventional GEM. Furthermore, we developed a method to rank genes based on newly collected data by leveraging the concept of unknomics. The findings of this study are expected to contribute to the efficient design of research using GE. However, because the information extracted by the LLM and the gene rankings may contain errors, it is essential for users to manually verify the data as a final step.

## SUPPLEMENTARY DATA

- SUPPL1_168genes.xlsx
- SUPPL2_evaluation_data.xls
- SUPPL3_gpt4o_llm_results_240802.jsonl
- SUPPL4_gpt4o_mini_llm_results_240802.jsonl
- SUPPL5_eval_results_targetedgenes_gpt4o.xls
- SUPPL6_eval_results_deg_gpt4o.xls
- SUPPL7_scores.xls
- SUUPL8_scores_modified_weights.xls

## ACKNOWLEDGEMENTS

This research was supported by the Center of Innovation for Bio-Digital Transformation (BioDX), the open innovation platform for industry-academia co-creation (COI-NEXT), the Japan Science and Technology Agency (JST; Grant Number JPMJPF2010).

This work was also supported by the JST, which established university fellowships for the creation of science and technology innovation (Grant Number JPMJFS2129). Computations were performed on the computers at Hiroshima University Genome Editing Innovation Center. We also would like to thank all laboratory members at Hiroshima University and the Database Center of Life Science (DBCLS) for their valuable comments.

## DATA AVAILABILITY

The Supplementary File and the source codes for this research are available at https://github.com/szktkyk/extract_geinfo, https://github.com/szktkyk/visualize_geinfo, and figshare repository (https://doi.org/10.6084/m9.figshare.c.7497327).

### Contributions

T.S. was responsible for the data curation, software development, pipeline analysis, draft of the original manuscript. T.S and H. B were responsible for the study design, conceptualization, methodology manuscript review and editing. H.B. was responsible for the project administration, funding acquisition. All the authors have read and approved the final version of the manuscript.

## ETHICS DECLARATIONS

Competing interests

The authors declare no competing interests.

